# RAMBO: Resolving Amplicons in Mixed Samples for Accurate DNA Barcoding with Oxford Nanopore

**DOI:** 10.64898/2025.12.29.694971

**Authors:** Andreas Kolter, Paul DN Hebert

## Abstract

DNA barcoding, the use of short genetic markers to identify and differentiate species, is a foundational tool for ecological and taxonomic research. The method has been scaled rapidly with next-generation sequencing technologies enabling the processing of thousands of specimens in parallel. Nanopore sequencing not only offers a flexible, low cost alternative to other platforms but produces full-length reads in real time and can be used in remote settings. However, its comparatively high error rate complicates downstream processing, particularly when PCR amplifies multiple templates from a single specimen, reflecting pseudogenes, paralogs, or contaminants. We present a novel pipeline for DNA barcoding that resolves mixed sequence signals from Nanopore reads using unsupervised clustering and staged consensus generation, without relying on curated reference databases, taxonomic priors, or error models. While existing methods to curate Nanopore sequence data assume a single dominant amplicon per sample or require deep sequence divergence among amplicons, our pipeline can distinguish variants differing by as little as 0.15 percent. It combines column-weighted encodings, UMAP projection, and HDBSCAN clustering, followed by conservative consensus refinement. The pipeline was benchmarked and validated using datasets with known composition, including high-fidelity PacBio sequences. The results show that Nanopore barcoding, when paired with appropriate analysis, can recover biologically meaningful variation even in technically complex samples. The pipeline is particularly suited for specimens where divergent templates are co-amplified, including mitochondrial pseudogenes or multicopy nuclear regions like ITS. As such, it provides a generalizable framework for high-resolution Nanopore analysis of complex amplicon mixtures.

## Introduction

Understanding global biodiversity is essential for tracking its loss, managing ecosystems, and informing conservation policy. Yet identifying species remains a major challenge in ecology, taxonomy, and environmental monitoring (Bickford et al., 2007; Goldberg et al., 2016; Li and Wiens, 2023; Löbl et al., 2023). Molecular approaches, particularly those employing DNA, offer scalable and replicable alternatives to traditional morphology-based approaches and have transformed how biodiversity is documented (Adams et al., 2023; Deiner et al., 2017; Goldberg et al., 2016; Pawlowski et al., 2021).

DNA barcoding identifies species by amplifying and sequencing standardized genetic markers, such as the mitochondrial cytochrome *c* oxidase subunit I (COI) gene in animals or the nuclear ribosomal internal transcribed spacer (ITS) region in fungi and plants from individual specimens (Chen et al., 2010; Hebert et al., 2003; Schoch et al., 2012). As Sanger sequencing provides sufficient read length for COI and ITS barcodes, it shaped the development of early protocols. However, it only produces clean data (chromatograms) when a single dominant PCR product is amplified, as co-amplified templates are typically masked or result in an entangled sequencing signal (Al-Shuhaib and Hashim, 2023; Kommedal et al., 2008).

Next-generation sequencing (NGS) platforms enable greater scalability as they produce millions of reads per sample (Porter and Hajibabaei, 2018), potentially enabling the separation of target amplicons from co-amplified templates (Shokralla et al., 2015). While short-read NGS platforms like Illumina are widely used for metabarcoding due to their high sequence output and low error rates, they cannot recover full-length 658 bp COI barcodes without assembly (Elbrecht and Leese, 2017; Shokralla et al., 2015). Although the Sequel II platform from Pacific Biosciences (PB) delivers highly accurate long reads, its high cost and large size restrict its use to core facilities (Wenger et al., 2019; Huo et al., 2021).

Sequencers from Oxford Nanopore Technologies (ONT) support full-length barcode recovery and their compact size and low cost makes them suitable for distributed workflows (Seah et al., 2020; Srivathsan et al., 2024; Hebert et al. 2025). However, their comparatively high error rate of 1–2.5% (R10.4, Dorado v0.31+), reflecting both miscalled bases and indels, particularly in homopolymer regions, requires the combination of several reads to generate an accurate consensus sequence (Seah et al., 2020).

Consensus sequence generation is effective when all reads from a sample originate from the same underlying PCR template, allowing errors to be averaged out to recover the true nucleotide sequence (Figure 1). This condition is met in DNA barcoding when a single-specimen sample features just one dominant amplicon (Srivathsan et al., 2024, Hebert et al. 2025). However, complexity is introduced when multiple templates within a sample are co-amplified (Figure 1). It can be resolved if the templates derive from distantly related taxa, such as organisms from a different family. In such cases, clustering approaches at 3% or higher can effectively identify all underlying templates, as the divergence between them exceeds the error rate of Nanopore by a substantial margin (Hebert et al., 2025). However, discriminating closely related sequences generated by co-amplification of mitochondrial pseudogenes, heteroplasmic variants, or contamination by congeneric species within a sample is challenging, as their sequence divergence may be less than or similar to the Nanopore error rate (Bensasson, 2001; Bonin et al., 2023; Hebert et al., 2023). These cases result in ambiguous consensus sequences or, more problematically, the merging of distinct biological entities into a single consensus that obscures the true biological signal, particularly when target mitochondrial sequences are conflated with ancestral pseudogenes (Parr et al., 2006; Song et al., 2008).

**Figure 1:**
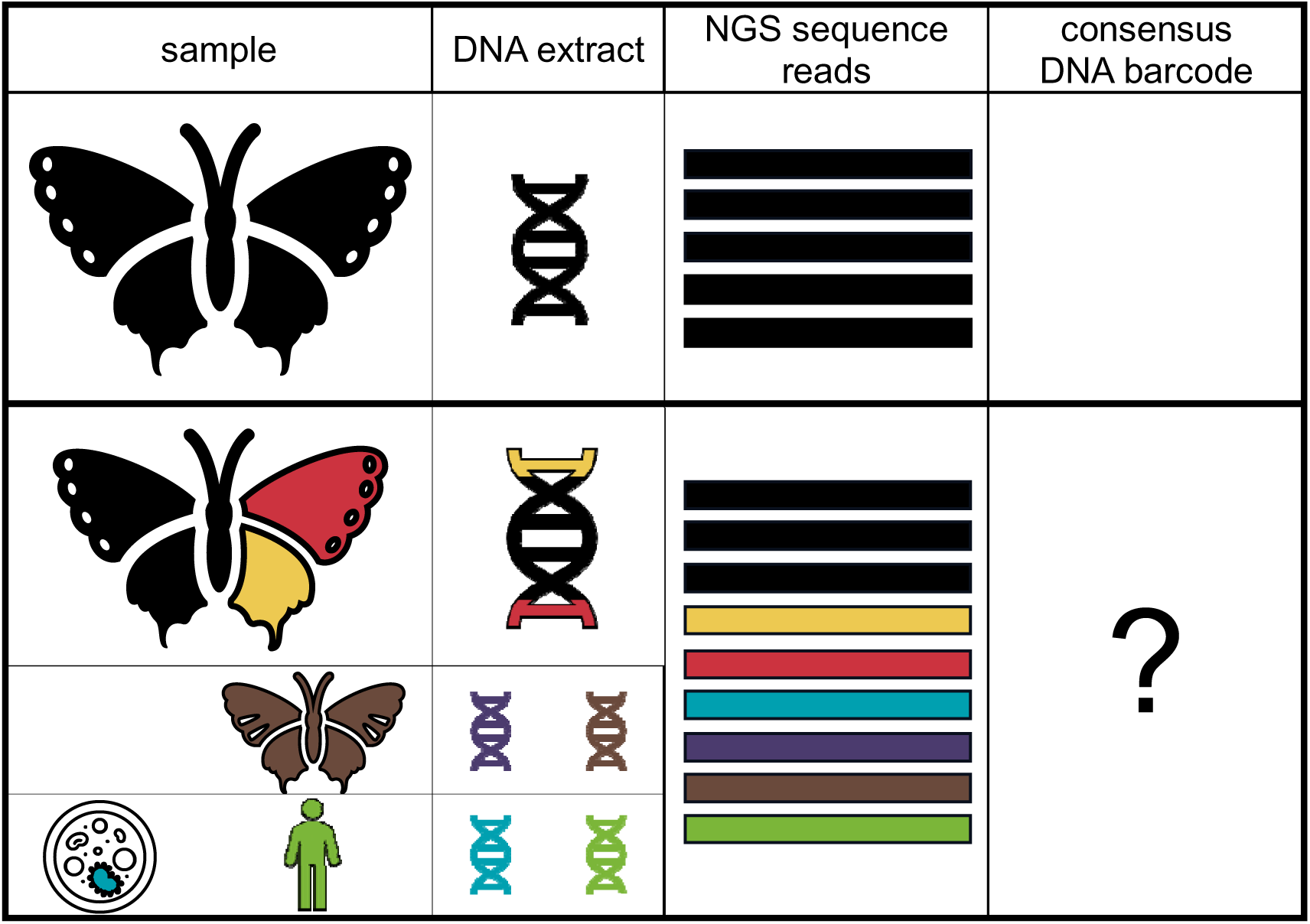
Row 1 shows an ideal barcoding case where DNA from a single specimen yields sequence reads from one template derived from the target organism, so consensus calling is straightforward. Row 2 shows complications of mitochondrial pseudogenes (yellow, red), which introduce two templates similar to the sample (black). Row 3 shows additional nontarget templates, including host associated ectoparasitic mites (purple) and microbes or endosymbionts (blue), as well as external contamination from other species (brown) and humans (green). All these templates can be co-amplified and sequenced, complicating consensus calling.

Computational strategies can address the limitations of Nanopore sequencing for mixed-template amplicon pools (Karst et al., 2021; Rodríguez-Pérez et al., 2021; Sahlin et al., 2021; Srivathsan et al., 2024). Reference guided genomic analysis methods such as variant calling and phasing infer variation relative to a reference genome, but they can be affected by reference bias when the reference is not representative. In biodiversity applications, comprehensive reference resources are often unavailable across taxa and regions, which limits the applicability and interpretability of reference-based analyses (Zhang et al., 2010). Read-to-read self-correction methods likewise assume long molecules with deep coverage, typically over 10 kilobases in length (Y. Liu et al., 2024), and are not applicable to barcode-length fragments. ONTbarcoder is tuned for short amplicons, but assumes a single dominant sequence per sample, an assumption that is violated in mixed-template contexts (Srivathsan et al., 2024).

To address the lack of methods tailored to mixed-template Nanopore amplicons at barcode length, we developed RAMBO (Resolving Amplicons in Mixed Samples for Accurate DNA Barcoding with Oxford Nanopore), a resolution-aware denoising pipeline for Nanopore amplicon data. RAMBO separates and denoises closely related sequences directly from the read set, without relying on reference genomes, genome assemblies, long-read self-correction, single-haplotype assumptions, externally trained error models, or curated taxonomic databases. It operates on demultiplexed raw reads and produces denoised sequence variants upstream of, and fully decoupled from, any taxonomic assignment. Using high-fidelity PacBio amplicons as ground truth, we validate its performance and quantify its resolution as the minimum sequence divergence at which variants remain reliably distinguishable in Nanopore read data.

The computational approach of combining UMAP (McInnes et al., 2018) with HDBSCAN (McInnes et al., 2017) builds on earlier pipelines (Rodríguez-Pérez et al., 2021). Additionally, the parallel development of PIKE, which also employs a closely related UMAP–HDBSCAN strategy, provides convergent support for the suitability of this approach to Nanopore amplicon clustering (Krivonos et al., 2025). RAMBO is implemented in R and integrates into existing bioinformatics workflows. We evaluated its performance on three datasets. The first contained pooled COI amplicons from multiple specimens of the same species and was designed to test if low (<1%) sequence divergence can be resolved. The second test examined COI barcode reads from a published study in which consensus calling failed. We re-analyzed these data with RAMBO to resolve base-call ambiguities and recover the correct barcode in the presence of extensive co-amplification. The third dataset consisted of nuclear ribosomal DNA amplicons from Euglossini bees (ITS1, 5.8S, ITS2, and part of 26S nrDNA) that had been validated against high-fidelity PacBio reference sequences. Together, these datasets show that RAMBO resolves sequence diversity within barcoding markers even when mixed templates are co-amplified.

## Materials and Methods

### Overview and Implementation

RAMBO is implemented as a R workflow for clustering and denoising by consensus generation of Nanopore amplicon reads without reliance on reference sequences. It produces one or multiple consensus sequences from mixed-input amplicon samples where each sample is represented by a FASTQ file. Performance-critical steps such as pairwise Hamming calculations were accelerated using custom C++ functions via Rcpp (Eddelbuettel et al., 2008). The pipeline depends on publicly available packages including Biostrings, dbscan, dplyr, FNN, ggplot2, igraph, magrittr, plotly, ShortRead, stringr and uwot (H. Pagès, 2017; Hahsler et al., 2019; Wickham et al., 2014; Beygelzimer et al., 2010; Wickham, 2009; Csárdi et al., 2025; Bache and Wickham, 2014; Sievert, 2020; Morgan et al., 2009; Wickham, 2016; Melville, 2019; Corporation and Weston, 2011; Microsoft and Weston, 2009). Throughout the pipeline, read-level quality was expressed as a log10 based Phred score corresponding to the mean per base error probability.

In short, RAMBO performs the following steps:

(1) It accepts primer-trimmed, demultiplexed, unaligned FASTQ reads and aligns them using MAFFT in FFT-NS-2 mode (Figure 2), preserving the quality scores. All downstream calculations operate on these exact aligned strings with no column rearrangements.

**Figure 2:**
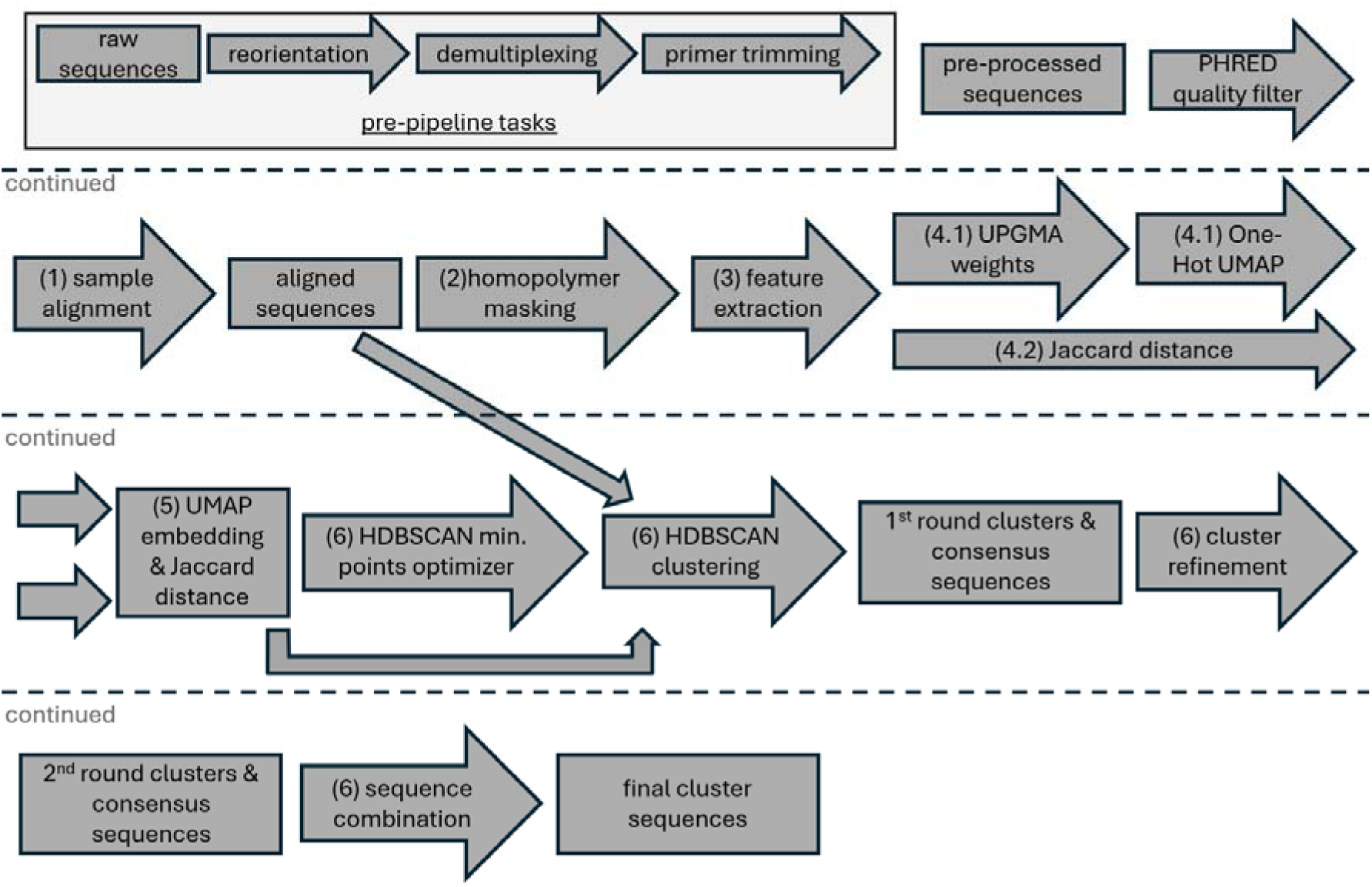
Overview of the RAMBO pipeline. Starting from preprocessed reads, the pipeline applies quality filtering, per sample alignment, homopolymer aware distance estimation, density-based clustering, consensus sequence calling with IUPAC ambiguity codes, and a clustering refinement to produce final cluster consensuses and metrics. Numbered elements in the schematic correspond to the bracketed step numbers in the Methods.

(2) Because homopolymer regions (default: >5 mono- or di-nucleotides) are susceptible to sequencing errors, they are masked to prevent technical artifacts from distorting downstream clustering. Homopolymers are restored after the clustering step, prior to consensus calling.

Homopolymer free sequence alignments were processed in four main stages: feature detection and feature encoding (3), distance calculations (4), distance adaptive merging (5) and density clustering (6):

(3) Informative features were defined at the level of individual alignment columns. For each column, every distinct nonconsensus base (A, C, G or T) that satisfied joint thresholds on minimum count and minimum frequency was treated as a candidate feature. To control false positives, candidate features were evaluated with a one-sided binomial test against a smoothed background error estimate. Features passing this test were encoded as binary presence or absence across reads.

(4) Based on these features, two distance metrics were constructed:

(4.1) First, a pairwise Hamming distance matrix was computed across all alignment columns containing features and subjected to hierarchical clustering with average linkage (UPGMA) using the R function hclust, yielding a dendrogram. This dendrogram was cut at a series of unevenly spaced heights, with increased sampling in regions where many merges occurred. Each cut partitioned the sequences into groups corresponding to the branches present at that height. Groups passing the filtering thresholds (default: >10 sequences and >1% of the total sample sequence count) were analyzed independently. For each clade, nucleotide frequencies for each column were computed for the clade versus all other sequences in the sample. To quantify differences in nucleotide composition between the clade and the remainder of the sample at each column, we used total variation distance, calculated as half the sum of the absolute differences between the frequencies of A, C, G, and T in the two groups. Columns with a total variation distance above a preset threshold (default 0.25) were marked as informative and assigned a score equal to that value. Because this procedure was repeated across all clades and cut heights, we applied a diminishing returns scheme in which the incremental weight assigned to a column at each additional hit was divided by the number of times that column had previously been flagged as informative. This prevented early, deep splits in the dendrogram from dominating the final column weights relative to finer scale differences near the tips. The column weights were applied in a one-hot encoding matrix, where each nucleotide at each alignment position was represented by a value which was the product of its Phred derived confidence (%) and the corresponding column weight. UMAP was then used to reduce the weighted feature matrix to five dimensions, so that each read was represented by five coordinates, summarizing its similarity to other reads across the original nucleotide-by-position features.

(4.2) Second, read to read distances were calculated by adaptively combining two complementary distance metrics derived from the binary feature matrix. (a) One metric used a Jaccard approach in which the weighted overlap of shared features (intersection) and the weighted set of features present in at least one read (union) were converted into a similarity score that emphasized shared minor allele patterns while downweighting features that were common across many reads. Because many read pairs contained only few detected features, joint absence across the retained feature set was additionally incorporated so that comparisons were not driven solely by a very small number of present features. (b) The second metric used a signed cosine distance calculated from a two-hot encoding of feature presence and feature absence. It distinguished read pairs with genuinely opposing feature patterns from pairs that were simply weakly informative because both contained little signal. The final read to read distance was obtained by adaptively blending metric (a) and metric (b), with mixing weights determined by per pair feature overlap and per read feature coverage.

(5) The two distance measures were combined into a single mixed distance that interpolates between Jaccard and UMAP based geometry. Jaccard distances on the binary feature matrix and Euclidean distances in UMAP space were each divided by their median over all finite entries, so that typical distances had similar numerical magnitude and neither component dominated purely by scale. For each read, an adaptive Jaccard weight was derived from its feature coverage: reads with few informative features received a higher Jaccard weight so that rare markers contributed more strongly, whereas reads with many informative features received a lower Jaccard weight and a correspondingly higher UMAP weight to suppress bridge forming dense feature patterns. For any read pair, the effective Jaccard weight was taken as the larger of the two read specific weights, the UMAP weight as the complementary fraction with a small minimum contribution, and the mixed distance was defined as the weighted average of the rescaled Jaccard and UMAP distances.

(6) Density based hierarchical clustering (HDBSCAN) was applied to the mixed distance matrix from the homopolymer-free masked alignments while scanning a grid of candidate minPts, where minPts defines the local neighborhood size used by HDBSCAN to estimate density and identify clusters. For each candidate minPts, a single clustering run yielded four metrics: the fraction of reads in non-noise clusters, the mean membership probability, a parsimony score reflecting the gap between the minimal allowed and largest observed cluster size, and an ambiguity penalty from the number of alignment columns with substantial secondary nucleotides within clusters. These metrics were rescaled to the interval [0,1] across candidates and combined by a fixed weighted sum favoring low noise, high membership probability, low ambiguity and moderate parsimony, and the neighborhood size with the highest score was retained. A final HDBSCAN run with this setting on the same mixed distance provided definitive cluster labels and outlier scores; reads assigned to the noise cluster or exceeding a fixed outlier threshold were flagged as outliers. For each remaining cluster, an IUPAC-aware consensus sequence (IUPAC threshold default: 25%) was computed from the non-homopolymer masked alignment, and clusters whose consensus sequences differed by at most a user defined Hamming distance were merged (default threshold: 0, implying no post hoc merging). Optionally, a UMAP implementation limited to three dimensions is capable of generating an interactive 3D cluster map (Supplementary Material 1a).

### Code and data availability

The full pipeline, including installation instructions, test data, and parameter documentation, is publicly available at https://github.com/Andreas-Bio/RAMBO. Benchmark datasets are deposited under 10.5281/zenodo.17940933.

### Dataset 1: Processing of Low-Divergence Barcode Samples

The first test dataset consisted of sequences recovered from 23 *Phyllocnistis populiella* (Lepidoptera, BIN BOLD:ACX5473) specimens represented in "Barcode 100K Specimens: In a Single Nanopore Run" (Hebert et al., 2025). These samples were chosen because their COI consensus sequences differed by between 1 and 10 nucleotides in pairwise Hamming distance, corresponding to 0.15% to 1.5% divergence, with a median of 4 differences (Supplementary Material 1b). This narrow divergence range was used to assess the pipeline’s ability to resolve fine-scale sequence variation.

Raw sequencing reads and consensus sequences were obtained from the Dryad repository (Prosser et al., 2024). Basecalling was performed using Dorado v0.3.1 in FAST mode, yielding reads with an average estimated error rate of 2.2%, corresponding to a Phred score of 16.6 (Supplementary Material 1c). Reads were all oriented into 5′–3′ direction and subjected to demultiplexing, primer trimming, and length filtering to retain fragments between 650 and 665 base pairs. Read counts per sample after filtering ranged from 90 to 379 with a median of 231 and a standard deviation of 57 (Supplementary Material 1d). Demultiplexing information was removed from the sequences and embedded in the read headers, and the 23 sample-specific read sets were pooled into a single FASTQ file for downstream analysis. Sample identifiers were retained from the original publication.

The denoising pipeline was executed using standard parameters. The average Phred quality threshold for alignment was set to 16 to accommodate the lower accuracy of FAST basecalls.

Finally, the same dataset has been analyzed with the PIKE pipeline in single mode, adjusting the maximum sequence length filter to 700, and the results were compared (Krivonos et al., 2025).

### Dataset 2: Resolving Mixed Templates to Remove Consensus Ambiguities in Nanopore Reads

The second test dataset consists of sequences from 66 insects (Supplementary Material 1e): 42 Diptera, 13 Hymenoptera and 11 belonging to other orders. The most abundant families were Phoridae (Diptera, 14 specimens) and Sciaridae (Diptera, 11 specimens). Samples were selected from the same study as dataset 1 based on their high number of ambiguous nucleotides (N >= 8) in the published COI barcode consensus sequence (Hebert et al., 2025). Raw ONT reads for these samples were reprocessed with RAMBO to evaluate its ability to recover unambiguous consensus sequences from challenging input. The RAMBO consensus calling used the same IUPAC threshold of 25%. These 66 samples comprised 9,519 raw reads in total, with read depths ranging from 89 to 243 (Supplementary Material 1e). Basecalling was performed using Dorado v0.3.1 in high-accuracy (HAC) mode, resulting in an average estimated read error rate of 1.8%. All remaining preprocessing steps and pipeline parameters were identical to those applied to the *Phyllocnistis* dataset.

### Dataset 3: Resolving Intragenomic ITS Variants: A Strategy for Accurate Marker Recovery from Multicopy Ribosomal Repeats

The third dataset was generated for this study and comprises nrDNA sequences of *Euglossini* (Apidae, Hymenoptera), more specifically the full ITS1, 5.8S, ITS2, and part of the 28S (approximately the first 2050 bp) nrDNA, totaling more than 5000 bp. First, samples were collected between 500 masl (meters above sea level) and 1440 masl in Kosñipata, Cusco, Peru (Supplementary Material 1f). One hind leg per specimen was exported to the Centre of Biodiversity Genomics, University of Guelph, Canada for DNA extraction, PCR and sequencing. PCR reactions employed GXL polymerase (Takara Bio Inc., Japan; Cat. No. R050A) and were performed in two rounds: 1) targeted template amplification and 2) indexing (Supplementary Material 1g & 1h). After pooling the indexed PCR reactions, each pool was sequenced on both a Nanopore PromethION flowcell and a PacBio Sequel II SMRT 8M flowcell, following standard protocols with negative size selection of 3 kb and below during library preparation. Preprocessing involved demultiplexing, primer trimming, and removal of a solllfar undescribed repetitive ITS1 region using Tandem Repeats Finder (Benson, 1999), to avoid artefactual inflation of genetic distances caused by uncharacterized sequence patterns (Supplementary Material 1i). Due to the complex pre-processing operations, all steps were performed in FASTA format. After preprocessing, samples were filtered by read count, retaining 369 paired samples (738 total), each with more than 100 reads from both PacBio (PB) and Oxford Nanopore Technologies (ONT) data. For samples with more than 500 reads, a random subset of 500 was used, and when read counts differed between technologies, the larger set was downsampled to match the smaller. Reads shorter than 5,000 bp were excluded. The total number of reads for each set of 369lllsamples was 135,703. The median read depth was 436 for both technologies, with median read lengths of 5,614 bp (ONT) and 5,639 bp (PB). The pipeline was run with default parameters, except that FASTA files were used as input.

## Results

### Dataset 1: Benchmarking Fine-Scale Clustering on Closely Related Barcode Sequences

To evaluate RAMBO’s performance on a realistic but tightly constrained dataset, we analyzed a *Phyllocnistis populiella* dataset (Hebert et al., 2025). A total of 5,463 reads were provided as input to the pipeline. After the initial low-quality filter, 4,937 reads remained (-9.6%). Subsequent alignment-based filtering used an average Phred score threshold of 16, further reducing the dataset to 3,351 reads (-32.1%). Of these, 3,096 (92.3%) were assigned to clusters, while 255 (7.6%) were classified as noise by the clustering algorithm and were excluded from further interpretation.

Each of the 23 samples was associated with a single dominant cluster (Figure 3). Dominant cluster sizes ranged from 60 to 214 reads, with a median of 132. Cluster purity, defined as the proportion of reads contributed by the dominant sample, ranged from 97.8% to 100%, with a median of 100%, indicating that clusters were effectively sample specific (Figure 3). The elevated proportions (> 40%) of noisy reads in BIOUG94608-F10 and BIOUG94608-C06 reflect reads that are distributed across many low-frequency sequence variants that are too dissimilar from one another to concentrate into a single dense region. This is consistent with a significantly higher within-group spread of the noise category in the distance space compared with the two retained clusters (betadisper F = 13.001, p = 5.39e-06; permutation p = 0.001). Subsampling down to 600 reads (∼18% of 3351) retained the pattern with each of the 23 samples possessing a unique cluster (Supplementary Material 1j).

**Figure 3:**
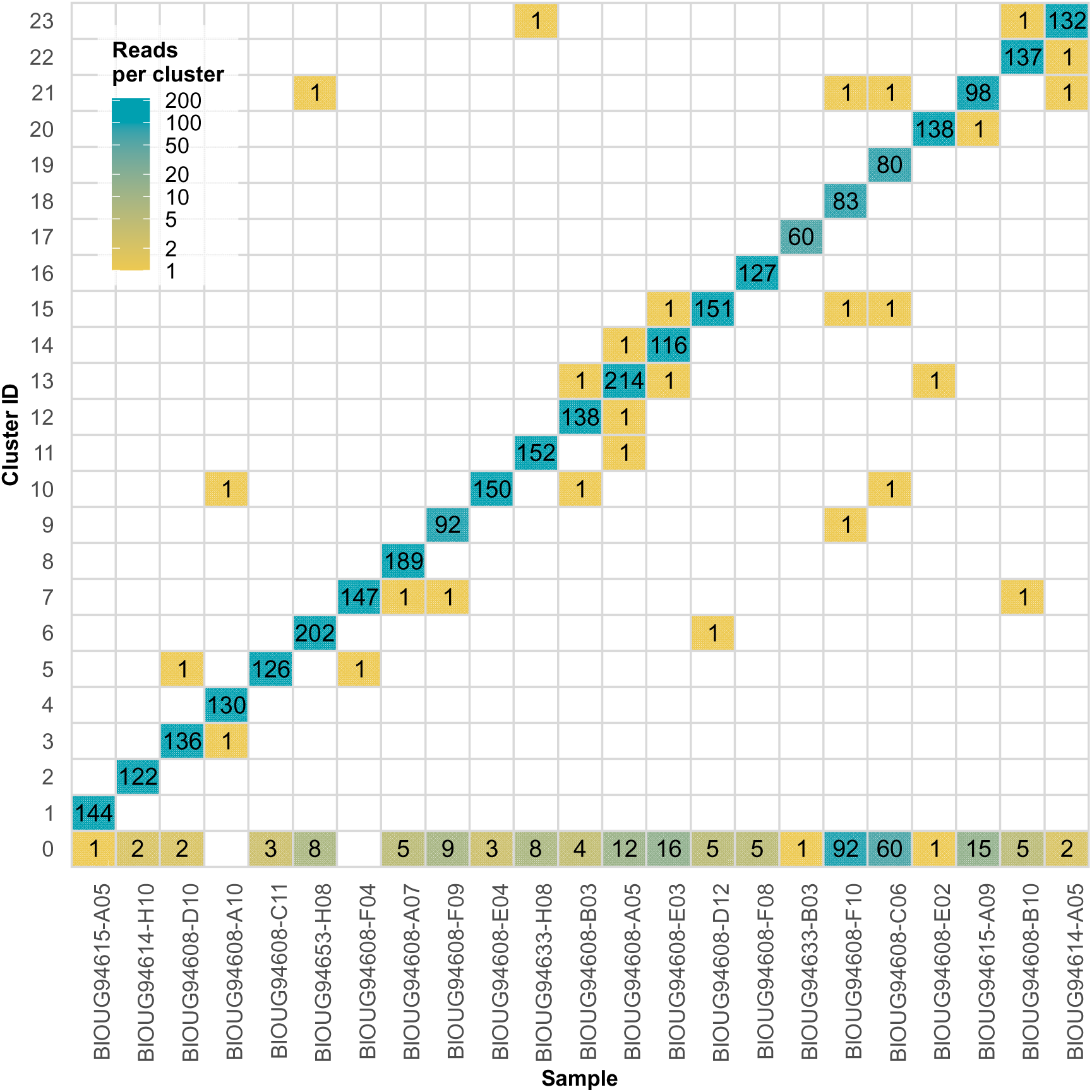
Sample-to-cluster assignments in dataset 1 (*Phyllocnistis populiella*) The 23 *Phyllocnistis populiella* samples (x-axis) are each assigned to one cluster (y-axis) by the RAMBO pipeline. Reads designated as noise (placed in Cluster ID 0) by HDBSCAN generally comprise a small fraction of the total reads per sample.

By comparison, the PIKE pipeline returned 59 raw clusters and 41 after size filtering with an average cluster purity of 66% (Supplementary Material 1k). Most notably, 12 of the 23 BIOUG samples were found in mixed clusters, largely because clusters whose consensus sequences differed by only 1 or 2 Hamming mismatches were not clearly separated.

### Dataset 2: Accurate Separation of Co-Amplified Sequences in Problematic Barcoding Cases

The 66 samples, selected for their high barcoding difficulty from the 100K dataset (Hebert et al., 2025), initially featured 686 Ns, while after processing with the RAMBO pipeline only 17 Ns (-97.5%) remained in consensus sequences (Figure 4). The median number of Ns dropped from 10 to 0 (Figure 4). The median number of reads included in the consensus decreased from 141 to 61 (Supplementary Material 1e). The RAMBO dominant cluster consensus had a similarity of 99.98% to previously published results when ambiguous nucleotides were interpreted based on IUPAC definitions (Supplementary Material 1e). Manual inspection of the non-dominant clusters revealed highly variable COI sequences to be the root cause of the ambiguous consensus.

**Figure 4:**
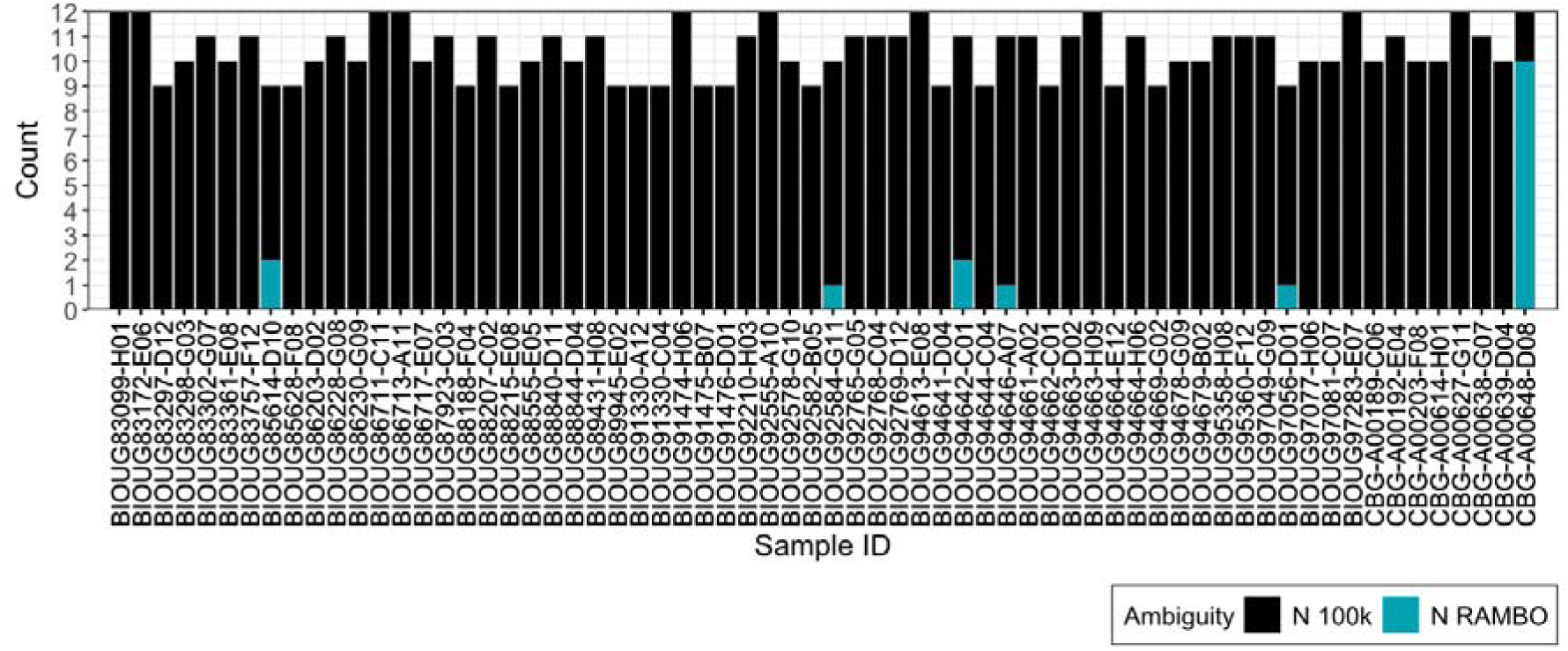
Comparison of Ambiguous Base Counts: 100K Study vs. RAMBO Output. Barplot illustrating the number of ambiguous bases in consensus sequences from the 100K study by Hebert *et al*. (2025) in black and the same raw reads reprocessed with RAMBO (cyan). Each bar represents a different sample with the y-axis showing the count of ambiguous nucleotides.

### Dataset 3: Multicopy nrDNA Marker Recovery across ONT and PacBio Platforms

We processed 738 samples in parallel (15 threads) in 407 min on an AMD Ryzen 5900X using between 32 GB and 64 GB RAM. Initial alignment lengths ranged from 5,728–10,682 bp (median 7,408 bp), consistent with expected insertions.

To test robustness of barcode consensus inference across sequencing platforms, we compared consensus sequences generated from ONT and PacBio reads of the same PCR amplicon pools, processed identically. PacBio was treated as a high-accuracy reference due to its read quality close to Q40. For each sample, instead of a strict comparison between the dominant ONT and PacBio cluster, candidate clusters for cross-platform comparison were selected from the two most abundant clusters per sequencing technology based on read count to avoid overinterpreting platform-dependent fluctuations in cluster dominance among closely competing clusters. The ONT–PacBio cluster pair with the highest sequence concordance was selected as the corresponding cluster pair for cross-platform comparison.

The mean sequence identity between results from the two platforms was 99.98% (Figure 5). Differences were driven primarily by indels, with a median of 1.73 gap openings per dominant cluster and a mean of 0.14 substitutions per cluster, corresponding to roughly one substitution per 41,500 bp. When gap openings were ignored, the mean identity increased to 99.998% while manual inspection indicated indel hotspots in and around homopolymer (N >5) tracts. Within clusters, per-read divergence was higher for ONT than for PacBio (1.60% versus 0.18%), but ONT consensuses remained stable and accurate, corresponding to an effective accuracy near Q35 (raw reads: Q22) (Supplementary Material 1c). Ambiguity in the dominant consensus was low, with ONT averaging 0.27 ambiguous nucleotides per sequence, about one per 21,000 bp, versus 0.18 for PacBio, about one per 33,000 bp (Figure 5). Median read support for dominant clusters was 115 reads for ONT and 108 for PacBio, and dominant consensus lengths had a median of 5632 bp with a range from 5353 to 6291 bp.

**Figure 5:**
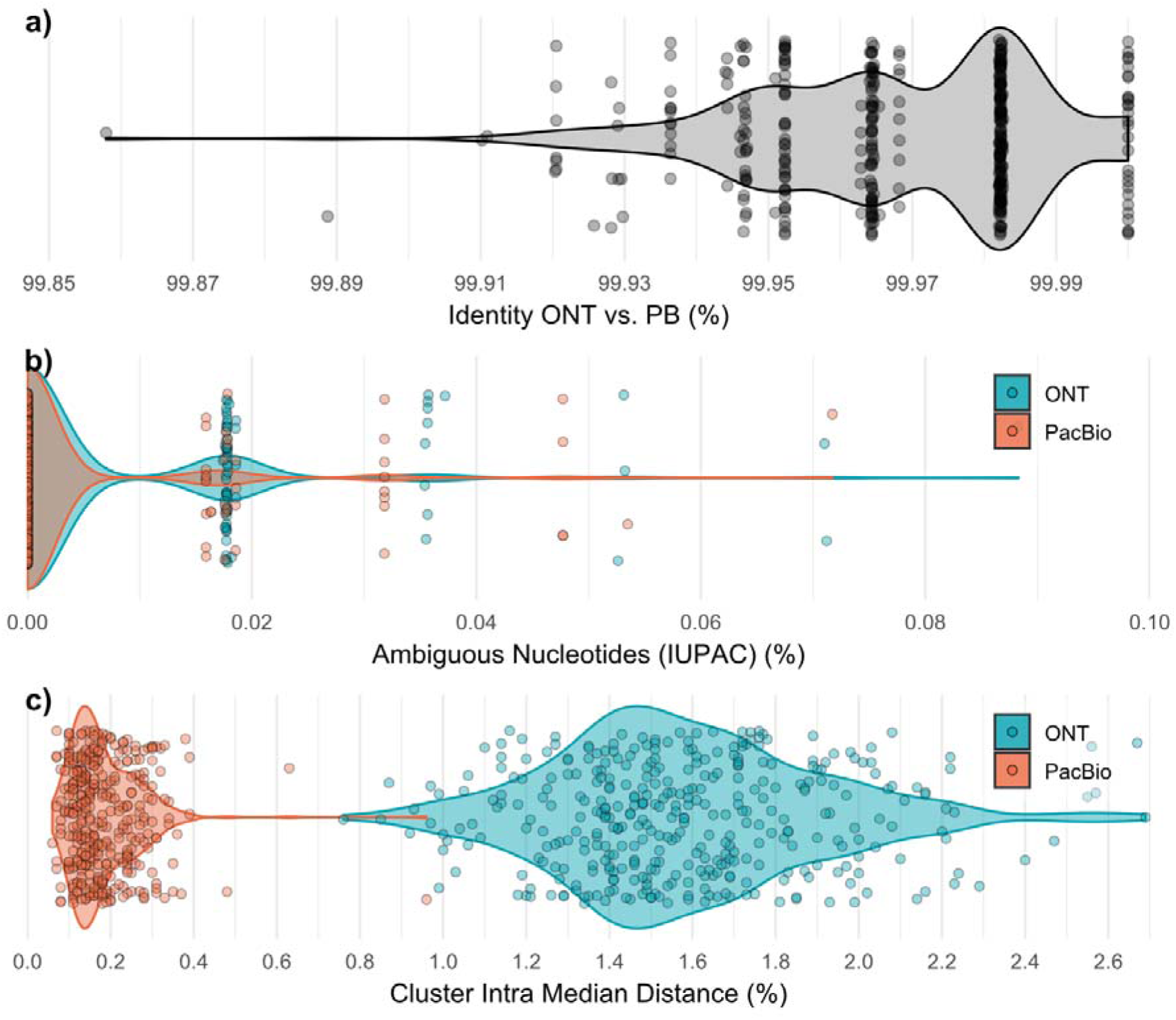
Euglossini ITS sequenced by ONT vs. PB. Violin plots show kernel density with width scaled by count. Points represent individual cluster values. a) Sequence identity between the consensus of the dominant ONT cluster and the consensus of the dominant PacBio cluster from the same specimen. Identity was computed on aligned sequences with IUPAC aware matching and terminal gaps ignored. The banded pattern is caused by discrete mismatch values (N=0,1,2,3…), offset by varying sequence lengths. b) Proportion of ambiguous nucleotides IUPAC in the ONT and PacBio consensuses of dominant clusters. Two outliers were removed for readability. c) Intra cluster median distance for ONT and PacBio, calculated as the median pairwise Hamming distance among the inlier reads for each cluster. Higher values indicate greater within cluster divergence. One outlier was removed for readability.

When we extended analysis to all clusters across 369 paired samples, PacBio produced more clusters per sample, mainly in the low-support regime. ONT yielded 1371 clusters in total with a median of three clusters per sample, whereas PacBio yielded 2111 clusters with a median of five clusters per sample. The paired difference in total cluster counts (ONT − PacBio) had a mean of −2.005 (Wilcoxon signed-rank V = 10,307, p = 1.8 × 10⁻¹[). For well supported clusters with at least 50 reads, ONT showed slightly more clusters per sample than PacBio, with a mean paired difference of +0.154 and a median of 0 (V = 10,478, p = 3.6 × 10⁻³). For clusters with fewer than 50 reads, PacBio produced more clusters, with a mean paired difference of −2.16 (V = 10,766, p = 1.5 × 10⁻¹[). Across all clusters, ambiguity remained low but the mean number of ambiguous IUPAC positions was higher for ONT than for PacBio (1.58 versus 0.32 per consensus), while the median was 0 for both. High-ambiguity clusters with more than 0.5% IUPAC positions were rare: we only observed 15 for ONT and none for PacBio, with median read depths of 23.

## Discussion

Reliable species identification depends on transparent, regimented data processing using protocols which ensure that observed sequence diversity reflects biological reality rather than methodological artifacts. For regulatory bodies and conservation practitioners, reproducibility and clearly defined performance thresholds are necessary before molecular methods can be integrated into monitoring frameworks. Without rigorous benchmarking and resolution metrics, it remains unclear whether a pipeline can distinguish closely related taxa or deliver consistent results across sample types.

### Dataset 1: Clustering Accuracy, Variant Recovery, and Pseudogene Detection

The fact that error-prone Nanopore reads complicate the discrimination of closely related taxa is well established, particularly when their sequence divergence approaches the scale of sequencing error (Pomerantz et al., 2022; Ohta et al., 2023). To mitigate this limitation, earlier approaches have often relied on prior knowledge of the expected amplicon sequence, requiring access to a high-fidelity reference sequence that can be used to model Nanopore error patterns during consensus generation (Espada et al., 2022). While effective in settings targeting one or a few well-characterized species, these strategies do not directly address resolution limits in closely related, unknown samples, a frequent situation in biodiversity studies.

By contrast, RAMBO operates without prior knowledge of sample identity or composition as sample identities are not used during clustering or consensus generation. Ground truth information is used exclusively for downstream validation of inferred clusters, not for guiding their formation. A subset of 23 *P. populiella* specimens with low pairwise divergence was used to benchmark the resolution of the clustering and consensus framework. Despite the small pairwise genetic Hamming distances (1–10), the pipeline accurately recovered all of the original sample identities. Discrimination down to 0.15% sequence divergence demonstrates RAMBO‘s ability to retain fine-scale variation while minimizing both over splitting and unjustified cluster merging. This is more than an order of magnitude improvement from values in literature, such as 2% in a fungal metabarcoding study (Dierickx et al., 2024), 5% for Amplicon_sorter and "Barcode 100K Specimens" (Vierstraete and Braeckman, 2022; Prosser et al., 2024), or the default cluster setting of 10% for Natrix2 (Deep et al., 2023) or PRONAME (Dubois et al., 2024).

### Dataset 2: Improved Consensus Accuracy in Difficult Barcoding Cases

RAMBO generated a fully unambiguous consensus sequence for 60 of 66 challenging samples, while the number of ambiguous bases was reduced in the others. This outcome demonstrates the pipeline’s ability to isolate the dominant COI signal by separating divergent or noisy reads that would otherwise introduce ambiguity, as reflected by the reduced sequence count contributing to each final consensus.

In the original 100K Nanopore barcoding study (Prosser et al., 2024), contamination and pseudogenes were mitigated by splitting sequences with more than 5% divergence into separate contigs. The persistence of ambiguous nucleotides in the targeted samples implies that the reads responsible for the ambiguous calls in these cases diverged from the target sequence by less than 5%. RAMBO addresses this challenge by applying dynamic similarity-based clustering and selecting the most internally consistent cluster with the highest read count for consensus generation, thereby avoiding reliance on fixed divergence thresholds.

In practical terms, the present results suggest that high-throughput barcoding datasets contain a recoverable fraction of sequences previously discarded or flagged due to ambiguity, and that unsupervised read clustering provides an effective route to restoring them. This not only increases the yield of barcodes for ecological and taxonomic analyses but also opens the door to recovering accurate sequences in challenging cases involving co-amplified pseudogenes, particularly in taxa such as Orthoptera and Formicidae where conventional methods frequently fail (Song et al., 2014; Cristiano et al., 2014; X. Liu et al., 2024).

### Dataset 3: Cross-platform consistency in nrDNA marker recovery

Long-read barcoding has largely relied on PacBio because its circular consensus sequencing protocol yields very high read accuracy and comparatively low exposure to indels in long amplicons (Wenger et al., 2019). In fungi, sequencing the nrDNA operon has been established using PacBio with mention of the possibility to extend the workflow to Nanopore (Runnel et al., 2022). However, the authors also mentioned inherent problems, such as widespread contamination in older samples and described intragenomic polymorphisms below 2% divergence. As well, Nanopore nrDNA workflows lack mock communities composed of closely related taxa, leaving their sensitivity insufficiently characterized (Ohta et al., 2023).

We selected the nrDNA region rather than COI because it poses a more challenging benchmark due to its greater length and long homopolymer stretches which stress alignment, gap placement, and hence consensus rules. Additionally, the nrDNA array from single specimens often includes divergent intragenomic variants (Feliner and Rosselló, 2007). Homopolymer tracts were masked before clustering during distance calculation, but not during consensus calling. Differences between consensus sequences in ONT and PacBio are best explained by indel errors, consistent with literature (Liu-Wei et al., 2024).

Overall, PacBio provides a stringent reference, but ONT was found to generate comparable consensuses under the same pipeline. However, clusters with a low read count (< 20) should be viewed with caution, as some of their signal might be masked by noise. Alternative options to our approach are Unique Molecular Identifier workflows when experimental goals justify the substantial additional complexity and costs (Baloğlu et al., 2021).

### Comparison with PIKE and NanoCLUST

PIKE, NanoCLUST, and RAMBO all perform unsupervised clustering of Nanopore amplicon reads using UMAP followed by HDBSCAN as the central step of sequence inference. A direct performance comparison with NanoCLUST was not possible because this pipeline is incompatible with read lengths shorter than 1,000 bp owing to an external dependency reflecting its design for genome polishing (Rodríguez-Pérez et al., 2021). This limitation motivated development of the PIKE pipeline, which arose independently and in parallel to RAMBO and is included in this study for comparison (Krivonos et al., 2025).

PIKE and NanoCLUST represent each cluster by a single polished sequence generated using Medaka (Oxford Nanopore Technologies Ltd, 2025). However, RAMBO retains and reports intracluster sequence variation during refinement rather than collapsing clusters to a single Medaka-polished sequence. In this study, we show that the underlying assumption of one amplicon per cluster, which allows sequence collapse at the cluster level, does not hold for PIKE in datasets containing closely related sequences (see Results). This distinction is relevant for data interpretation, especially for applications in which single nucleotide differences are informative, such as plant barcoding, where individual SNPs can determine species-level assignment (Chen et al., 2013; Kolter and Gemeinholzer, 2021; Poliakova et al., 2020), as well as population-level analyses using COI (Matumba et al., 2020; Mohamadzade et al., 2023; Toli et al., 2023). Retaining information about heterogeneous clusters is pivotal to inform the user about cluster purity and to prevent the overinterpretation of results. It also allows intragenomic nucleotide variation of multicopy barcoding markers, such as variation commonly observed in telomere-proximal regions of nrDNA repeats (Chen et al., 2023), to be incorporated into the cluster signal without imposing prior assumptions about whether such variation is phylogenetically informative or not.

Further limitations concern how pipeline performance is evaluated. In both PIKE and NanoCLUST, performance is primarily assessed using mock communities and reported in terms of taxonomic completeness or correctness (Krivonos et al., 2025; Rodríguez-Pérez et al., 2021). Although mock communities are a valuable validation tool, taxonomy-based metrics conflate sequence inference with primer performance and matching logic, such as user-defined minimum coverage or maximum allowed divergence thresholds. Consequently, such evaluations do not provide a general benchmark for evaluating the performance of sequence reconstruction and do not allow direct assessment of the minimal divergence that can be discriminated. The only metric that directly reflects pipeline performance is the sequence identity of the inferred sequences relative to the ground truth, which is, in practice, a high-fidelity proxy, such as PacBio-derived sequences.

In summary, all three pipelines produce sequence data suitable for ecological applications. However, when closely related taxa must be resolved separately, caution is warranted because the method of consensus sequence generation determines how much informative variation is retained.

### Limitations

While other pipelines produce results from as few as five reads (Prosser et al., 2024), they presume the absence of secondary amplicons. The datasets analyzed in this study have a read depth of approximately 100 reads/sample. We tested the pipeline with samples between 20 and 10,000 reads per sample and downsampling is recommended for deeper libraries. Despite RAMBO‘s higher read requirement, the cost to obtain 100X read depth is minimal at scale. A PromethION flow cell produces an average of 100 million reads for $1000 US, corresponding to 1000 reads per $0.01.

The high resolution of RAMBO arises from its capacity to preserve site-specific informative positions throughout clustering. This strategy separates closely related variants but assumes that sequences are globally alignable and relies on pairwise alignment, which increases computational cost. For classical barcode libraries with broadly uniform reads, the resulting gap structure is well behaved and within design limits. By contrast, highly diverse mixtures with roughly equal contributions from multiple families are outside RAMBO’s intended use. In such cases, many alignment columns become dominated by gaps, including informative positions, and are removed by the global gap filter. This behavior matches metabarcoding scenarios rather than single specimen barcoding, and RAMBO was not designed for such use. The algorithm has been tuned for samples with up to about 10,000 reads.

RAMBO explicitly supports multicopy markers by preserving within-sample heterogeneity while limiting excessive fragmentation into small spurious clusters. Separate clusters are generated only when an alternative variant is supported by a coherent group of reads that concentrates into a compact high-density region in UMAP embedding. Reads forming diffuse, low-density clouds in the UMAP embedding are assigned to the noise class, consistent with the elevated noise fractions in BIOUG94608 F10 and BIOUG94608 C06 (Figure 3). Within well-formed clusters, residual variation that does not co-segregate into a separate dense read group is retained in the consensus as IUPAC ambiguity codes, protecting the integrity of multicopy markers such as ITS without inflating cluster counts. In cases where read depth is insufficient to resolve multiple distinct variants, several strongly supported sequence patterns can be absorbed into a single heterogeneous cluster. These clusters typically yield consensus sequences with elevated IUPAC ambiguity and can therefore be flagged and excluded.

### Conclusions and Outlook

Across the trial datasets, RAMBO met three demanding goals. First, it separated closely related sequences down to roughly 0.15% divergence while avoiding oversplitting, showing that fine-scale variation can be retained without fragmenting true biological units. Second, it recovered the dominant barcode from mixtures that contained co-amplified pseudogenes, so that these templates did not inflate ambiguity in the final sequence. Third, when coverage was adequate, it produced ONT consensus sequences that were highly concordant with their PacBio counterparts, indicating that Nanopore barcoding can deliver PacBio-level accuracy at the consensus level.

These results clarify where RAMBO sits within the growing ecosystem of tools for the analysis of amplicon pools sequenced on ONT. For COI barcoding taxa with no pseudogenes, ONTbarcoder is adequate because it assumes a single dominant mitochondrial haplotype per specimen (Srivathsan et al., 2024). When libraries combine strongly divergent barcoding markers in a single sample, Amplicon_sorter can perform well because the markers are separated by large sequence distance (Vierstraete and Braeckman, 2022). RAMBO has a different goal; it targets the more subtle cases where several closely related templates co-occur in a barcoding sample. These include nuclear mitochondrial pseudogenes, plastid paralogs co-amplifying with functional copies and multicopy nuclear markers such as ITS with genuine intra-individual heterogeneity. Within limits, RAMBO can tolerate and cluster co-amplified highly divergent reads from endophytic fungi, parasites or gut contents (Figure 1), if these divergent template sequences do not form the majority of reads. Large barcoding campaigns also introduce mixed individuals, low level contamination, and tag jumping. In all these situations, the reads form a community of closely and more distantly related templates rather than a single homogeneous amplicon, and RAMBO is designed to resolve that structure.

Looking ahead, RAMBO provides a path to high-resolution metabarcoding on ONT. A light front end that assigns reads into broad taxonomic groups (e.g., family) using reference matches, would separate deep divergences among lineages from the shallow variation within species that RAMBO is designed to resolve. Once reads are partitioned in this way, the same clustering and consensus strategy can be applied to mixed environmental samples while still distinguishing closely related species. Because family-level assignments are already reliable at Nanopore error rates, this extension is conceptually simple yet carries large practical impact. It would turn low cost long-read barcoding into a general metabarcoding tool that can follow fine-scale changes in community composition, unlike current Nanopore metabarcoding pipelines.

## Data Accessibility Statement and Acknowledgements

Scripts and data used to generate the results in this publication are available from: https://github.com/Andreas-Bio/RAMBO. All output files for all pipeline steps and the final cluster sequences can be downloaded at DOI: 10.5281/zenodo.17940933.

We gratefully acknowledge the Asociación para la Conservación de la Cuenca Amazónica (ACCA) and Estación Biológica Manu in Peru for logistical support and research access, especially the ICFC Canada Fellows for sample acquisition. Insects were collected and exported with permissions MIDAGRI-SERFOR-DGGSPFFS-DGSPFS: D000061-2022 & D000052-2023 & D000358-2023 & D000120-2024.

An AI-based language and coding assistance tool (ChatGPT 4.1) was used during manuscript preparation to support language editing and code review. All analytical decisions, code implementation, validation, and interpretation of results were performed and verified by the authors.

